# *Tomato spotted wilt virus* manipulates the reproduction of its insect vector, western flower thrips (*Frankliniella occidentalis*), to facilitate transmission

**DOI:** 10.1101/598920

**Authors:** Yanran Wan, Sabir Hussain, Baoyun Xu, Wen Xie, Shaoli Wang, Youjun Zhang, Xuguo Zhou, Qingjun Wu

## Abstract

**BACKGROUND:** *Tomato spotted wilt virus* (TSWV), one of the most devastating viruses of ornamental plants and vegetable crops worldwide, is transmitted by the western flower thrips, *Frankliniella occidentalis* (Pergande), in a persistent-propagative manner. How TSWV manipulates the reproduction of its vector to enhance transmission and whether infection with TSWV changes the mating behaviour of this thrips vector are not fully understood.

**RESULTS:** In this study, we found that TSWV-exposed thrips, in general, had a significantly longer developmental time than did non-exposed individuals. Such an increase was predominantly seen in adults, a stage associated with dispersal and virus transmission. TSWV-exposed *F. occidentali*s produced substantially more progeny than did non-exposed thrips. Interestingly, most of the increase in progeny came from an increase in males, a sex with a greater dispersal and virus transmission capability. Specifically, the sex ratio of progeny shifted from female biased (2-7:1) to evenly split or male biased. Regarding mating behaviour, compared to virus-free controls, TSWV-exposed *F. occidentali*s had significantly longer copulation duration, were more active in males, and remated less often in females.

**CONCLUSION:** These combined results suggest that TSWV alters the reproductive behaviour of its insect vector, *F. occidentalis*, to promote virus transmission. Consequently, a monitoring program capable of earlier detection of the virus and a reduced economic threshold for vector (thrips) control should be in consideration for the long-term, sustainable management of TSWV.

## 1 INTRODUCTION

The western flower thrips, *Frankliniella occidentalis* (Pergande), is an important agricultural pest with a worldwide distribution.^1^ *F. occidentalis* oviposits and feeds on vegetable, fruit and ornamental crops, causing substantial economic losses.^2-3^ In China, *F. occidentalis* was first detected in 2003 and has now become one of the most serious pests in greenhouse production.^4^

*F. occidentalis* has a haplodiploid reproductive system and can reproduce sexually or via arrhenotokous parthenogenesis, in which fertilized eggs develop into females and unfertilized eggs develop into males.^5^ In the mating system of *F. occidentalis*, females are relatively monandrous but usually can re-mate five days after the initial mating, and males are polygynous.^6^ In nature, the sex ratio of *F. occidentalis* is female biased, and the percentage of females ranges from 70 to 90%, i.e., female:male ratio of 7:3 to 9:1.^7,8^ However, the sex ratio of *F. occidentalis* can fluctuate under different biotic and abiotic conditions, including temperature^9^, host plant and season^3^, population density^7^, and pesticide treatment.^10^

Approximately 80% of plant viruses are transmitted by insect vectors.^11,12^ *Tomato spotted wilt virus* (TSWV) belongs to the genus *Tospovirus* and family Bunyaviridae and can infect more than 900 plant species in 90 plant families.^13^ As an emerging plant pathogen with global importance^14^, TSWV is transmitted exclusively by thrips.^15,16^ The virus must be acquired by the first instar for successful transmission, and adults are the primary plant-to-plant transmission stage because they are winged and highly mobile.^17^ *F. occidentalis* transmits TSWV in a persistent, propagative manner and has no transovarian transmission.^18^ TSWV transmission can be affected by host plants, virus load, age, sex, susceptibility to pesticide, and behaviour of thrips.^18-23^ Stafford et al.^22^ reported that carrying TSWV altered the feeding behaviour of *F. occidentalis*, in which exposed males fed significantly more than the un-exposed controls. In addition, exposure to TSWV increased the reproduction rate of *F. occidentalis*.^11,24,25^

*F. occidentalis* is a serious problem because of its feeding damage, and more importantly, its transmission of TSWV.^23^ Multiple lines of evidence suggest that plant viruses manipulate their vectors to enhance their transmission^22,26,27^, which has important implications in the ecology and evolution of pathogens. Wang et al.^28^ found that a novel negative-stranded RNA virus could mediate the secondary sex ratio of its host wasp by decreasing the percentage of female offspring, and because males transmit the virus to their offspring and can mate with multiple females, the virus would further spread. TSWV can replicate within its vector thrips but cannot transmit vertically^15,25^. How TSWV manipulates the reproduction of its vector to enhance transmission and whether infection with TSWV changes the mating behaviour of this thrips vector are not fully understood. To answer these questions, we investigated 1) the effects of TSWV on the development, longevity, and fecundity of *F. occidentalis* and 2) the effects of TSWV on the offspring of *F. occidentalis* using different parental sex ratios (3:1 or 1:1) and mating patterns (in one pair or paired in groups). Then, four cross-pair treatments were designed to determine which parent thrips (male or female) have been affected by TSWV and to study whether the mating behaviour of *F. occidentalis* is manipulated by TSWV. Our work shows that TSWV-exposed *F. occidentalis* produce significantly more males, resulting in the offspring sex ratio shifting from female biased to evenly split or male biased. TSWV-exposed *F. occidentali*s had significantly longer copulation duration, were more active in males, and re-mated less frequently in females. Our results suggest that TSWV alters the reproductive behaviour of its insect vector, *F. occidentalis*, to promote virus transmission.

## 2 MATERIALS AND METHODS

### 2.1 *Frankliniella occidentalis* strains and TSWV maintenance

Ivf03 is a laboratory susceptible strain and Spin-R is a spinosad-resistant strain as described by Wan et al.^4^ Ned, another laboratory susceptible strain, was a gift from Dr. Dick Peters, Wageningen University & Research, in 2006. All three strains were maintained separately on kidney bean pods at 26°C, 70% RH and a 16:8 (L:D) h photoperiod. TSWV isolate TSWV-YN was maintained on *Datura stramonium* plants by thrips transmission.^23^

### 2.2 TSWV on the development, longevity, and fecundity of *F. occidentalis*

Ivf03 and Spin-R strains were used in this experiment. Eggs of Ivf03 and Spin-R strains were obtained by placing fresh bean pods in glass jars containing hundreds of adults. Adult females were allowed to oviposit on the bean pods for 24 h. Thereafter, the adults were removed, and the bean pods with eggs were transferred to another jar, which was placed in an incubator at 26 ± 1°C with an L16:D8 h photoperiod until the nymphs hatched.

Mature leaves of healthy or TSWV-infected *D. stramonium* were collected, and a leaf disc (1 cm in diameter) was placed with its adaxial surface downwards on a sloping bed of agar (3 ml of 20 g/L) in a 5-ml centrifuge tube. Approximately 200 newly hatched Ivf03 and Spin-R nymphs (< 6 h old) were transferred individually into the each tube using a fine paint brush. A small hole (5 mm in diameter) in the lid of the centrifuge tube provided ventilation. Leaf discs were changed when necessary. The survival of each individual was recorded daily. When virus-exposed (fed on TSWV-infected *D. stramonium* during preadult stage) and non-exposed (fed on healthy *D. stramonium* during preadult stage) adults emerged, pairs were placed in plastic cylinders (15 cm high and 5 cm diameter) containing kidney bean pods (one female, one male, and one bean pod per cylinder). The bean pods were changed every two days. Each treatment (virus exposed vs. non-exposed for two strains of the thrips) was represented by 30 pairs of adults. The fecundity of females was recorded (determined from the number of the first instars^29^ that hatched on the replaced bean pods) until the females died. The replaced bean pods with eggs were placed in an incubator at 26 ± 1°C with an L16:D8 h photoperiod. The sex of each emerging adult was determined, and the numbers of females and males of each pair were counted. This experiment was replicated three times.

The developmental time, longevity and fecundity of virus-exposed vs. non-exposed parents of each strain were compared by one-way analysis of variance (ANOVA, *P*≤0.05). The percentage of males produced by the virus-exposed vs. non-exposed parents of each strain was compared by chi-square tests (*P*≤0.05).

### 2.3 TSWV on the offspring of *F. occidentalis* in group pairing

Newly hatched nymphs (< 6 h old) from Ivf03, Spin-R, and Ned were obtained as described in the previous experiment. A fine paint brush was used to transfer the nymphs to 15-cm-diameter Petri dishes (approximately 500 nymphs of one strain in each dish) containing pieces of healthy or TSWV-infected *D. stramonium* leaf on the moistened filter paper. The leaves were changed daily until adults emerged. Newly emerged females and males of each treatment were collected and placed in thrips-rearing jars containing kidney bean pods; each jar contained one bean pod and 45 females and 15 males or 45 females and 45 males. Each treatment ((three strains × ± virus exposure) × two ratios of female/male) was represented by three replicated jars. The bean pods were changed every two days. The replaced bean pods with eggs were placed in an incubator at 26 ± 1°C with an L16:D8 h photoperiod. When adults emerged, female and male thrips were counted.

The mean numbers of male, number of female and total fertilities of virus-exposed vs. non-exposed parents of each treatment were compared by one-way ANOVAs (*P*≤0.05). The percentage of males produced by the virus-exposed vs. non-exposed parents of each treatment was compared by chi-square tests (*P*≤0.05).

### 2.4 TSWV on the mating behaviour of *F*. occidentalis

#### Offspring in group cross pairing

Four cross-pairing treatments were designed as follows: Female (-) × Male (-) = non-exposed female × non-exposed male (treatment 1); Female (+) × Male (+) = virus exposed female × virus exposed male (treatment 2); Female (+) × Male (-) = virus exposed female × non-exposed male (treatment 3); and Female (-) × Male (+) = non-exposed female × virus exposed male (treatment 4). TSWV-exposed and non-exposed thrips were prepared as described in 2.3. For each treatment, freshly emerged adults of Ivf03 strain were collected and placed in thrips-rearing jars containing kidney bean pods and each jar contained one bean pod and 10 males and 10 females. Each treatment was replicated three times. The thrips maintained and offspring counted as described in 2.3. Differences in the average numbers of male and female of the four treatments were compared by one-way ANOVAs (*P*≤0.05).

#### Mating behaviour recording and offspring counting

Each cross-paired (as described above) virgin female and male (2 days old) of Ivf03 strain was video-taped using a Leica model DM 2500® microscope with a camera. A small arena (8 mm wide and 1.5 mm high) was made, and the thrips (one male and one female) were placed inside for observation. The virgin pairs were transferred carefully to the arena using a camel hair brush and were immediately covered with a glass cover slip. The videos were recorded for 1 h continuously for each cross-pairing treatment. Video playback recording was used to measure the pre-copulation (initial contact to male mounting and number of rejections by female prior to mating), copulation behaviour (duration, male stroking female, and calm period)^30^, and female mating frequency. Re-mating frequencies were observed in a 5-day interval^6^ for two times. Male harassment rates (repeated mating attempts by a male per hour) were also determined during the experiment.^30-32^ The TSWV acquisition by exposed male and female thrips was determined for each individual using RT-PCR^23^. The thrips that did not test positive for TSWV infection and that accidently killed were excluded from the replicates. Twenty replicates were included in all treatments.

On each recording day, mated females were removed after 1 h of recording and were transferred to 1.5-ml tubes containing a section of bean pod for 24 h to allow them to deposit eggs. The bean pods were changed and moved to an isolated petri dish for determination of the numbers of male and female offspring. The numbers of offspring per female were determined during its life for up to 20 days in all treatments.

Differences in the means for pre-copulation (log transformation), copulation duration (log transformation), female mating frequency, and male harassment rate among the four treatments were analysed using one-way ANOVAs (*P*≤0.05). The differences in the re-mating frequencies among four treatments were compared by chi-square tests (*P*≤0.05). The mean numbers of male and number of female per female of the four treatments were compared by one-way ANOVAs (*P*≤0.05).

### 2.6 Statistical analysis

Before each analysis, different measurements within each treatment repeat were tested for normal distribution (Shapiro-Wilk normality test at *P* > 0.05). In all analyses, the level of significance was set at α=0.05. SPSS software (version 19.0) was applied for analyses.

## 3 RESULTS

### 3.1 TSWV on the development, longevity, and fecundity of *F. occidentalis*

For both the Ivf03 and Spin-R strains, the developmental times of nymph and pupa were significantly shorter when fed on TSWV-infected than on healthy *D. stramonium* leaves (Fig. 1). In addition, the longevity of males and females was significantly longer for virus-exposed than non-exposed thrips, except for Spin-R males (Fig. 1). Fecundity was significantly higher for virus-exposed than non-exposed thrips in both strains (Table 1). Specifically, the male percentage was significantly higher in the progeny of virus-exposed than non-exposed thrips (Table 1).

**Table 1.** The parental fecundity and the percentage of males in the progeny of Ivf03 and Spin-R strains of *F. occidentalis* as determined by the TSWV status (virus exposed or non-exposed) of the parental thrips (one female and one male).

| Strain | TSWV status | Fecundity<br>(first instars/female) | Male<br>Percentage (%) | $\chi^2$ | *df* | *P* |
| --- | --- | --- | --- | --- | --- | --- |
| Ivf03 | Non-exposed | 141 ± 2 | 12 | 70.31 | 1 | <0.0001 |
| | Virus exposed | 255 ± 2* | 49 | | | |
| Spin-R | Non-exposed | 71 ± 3 | 21 | 171.95 | 1 | <0.0001 |
| | Virus exposed | 140 ± 1* | 49 | | | |

For each strain, a one-way ANOVA was used to determine the effect of virus exposure on the parental fecundity (an asterisk indicates a significant difference in fecundity at $P \leq 0.05$ ), and a chi-square test was used to determine the effect of virus on the percentage of males in the progeny.

**Figure 1.**
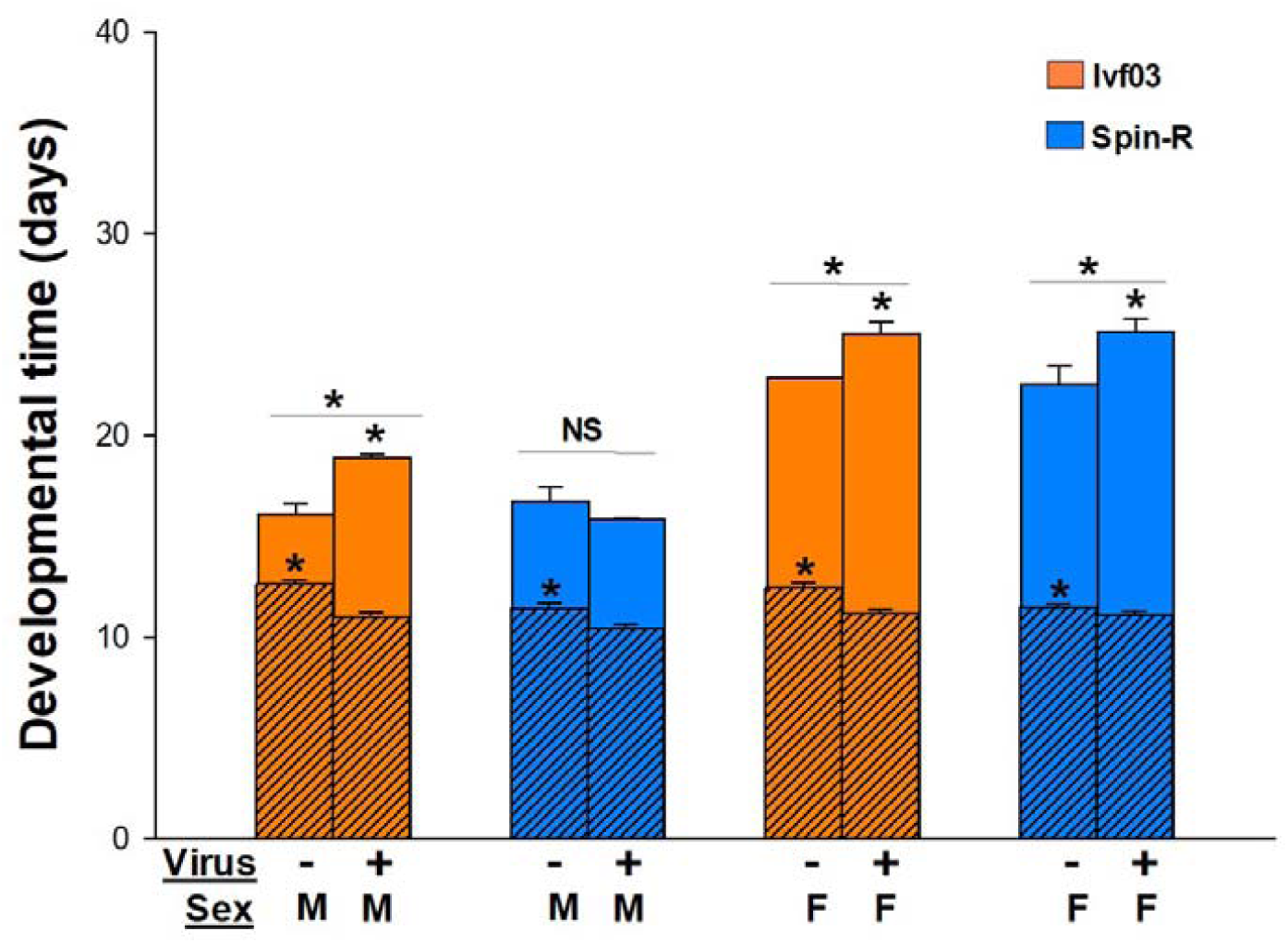
Developmental time of pre-adult (diagonal texture) and adult stages (blank texture) of Ivf03 and Spin-R strains of *F. occidentalis* as determined by the TSWV status (virus exposed or non-exposed) of the parental thrips. An asterisk (*) indicates a significant difference (one-way ANOVA, *P* ≤ 0.05). Values are means (± SE) of replicates. A minus sign (-) indicates non-exposed treatment and a plus sign (+) indicates virus exposed treatment. M represents male and F represents female.

### 3.2 TSWV on the offspring of *F. occidentalis*

Regardless of whether the parental sex ratio was 3:1 (45 females and 15 males) or 1:1 (45 females and 45 males), the total number of offspring per group was greater for virus-exposed parents than for non-exposed parents of all three strains (Fig. 2) When the TSWV-exposed parental sex ratio was 3:1, both male (2.8- to 5.2-fold) and female (1.9- to 2.3-fold) progenies were increased when exposed to TSWV. While under the 1:1 parental sex ratio, only the male progeny increased significantly (approximately 1.5-fold), and the number of female progeny in TSWV-exposed Ivf03 and Spin-R strains was even less than that in their corresponding non-exposed strains (Fig. 2). The percentage of males in the offspring was greater for virus-exposed parents than for non-exposed parents of all three strains regardless of the parental sex ratio (Tables 2 and 3).

**Table 2.** The percentage of males in the progeny of Ivf03, Spin-R, and Ned strains of *F. occidentalis* as determined by the status (virus exposed or non-exposed) of parental thrips (in groups of 45 females and 15 males).

| Strain | TSWV status | Male percentage ( % ) | | | | $\chi^2$ | df | P |
| --- | --- | --- | --- | --- | --- | --- | --- | --- |
|  |  | Rep.1 | Rep.2 | Rep.3 | Mean |  |  |  |
| Ivf03 | Non-exposed | 38 | 36 | 33 | 36 | 1017.81 | 1 | <0.0001 |
|  | Virus exposed | 68 | 61 | 64 | 64 |  |  |  |
| Spin-R | Non-exposed | 27 | 20 | 31 | 26 | 833.945 | 1 | <0.0001 |
|  | Virus exposed | 52 | 50 | 55 | 52 |  |  |  |
| Ned | Non-exposed | 34 | 38 | 38 | 37 | 515.11 | 1 | <0.0001 |
|  | Virus exposed | 55 | 51 | 55 | 54 |  |  |  |

Analyses were conducted in software SPSS (version 19.0) by chi-square analysis.

**Table 3.** The percentage of males in the progeny of Ivf03, Spin-R, and Ned strains of *F. occidentalis* as determined by the status (virus exposed or non-exposed) of parental thrips (in groups of 45 females and 45 males).

| Strain | TSWV status | Male percentage ( % ) | | | | $\chi^2$ | df | P |
| --- | --- | --- | --- | --- | --- | --- | --- | --- |
|  |  | Rep.1 | Rep.2 | Rep.3 | Mean |  |  |  |
| Ivf03 | non-exposed | 31 | 36 | 33 | 33 | 348.44 | 1 | <0.0001 |
|  | virus exposed | 48 | 54 | 51 | 51 |  |  |  |
| Spin-R | non-exposed | 27 | 35 | 32 | 32 | 548.22 | 1 | <0.0001 |
|  | virus exposed | 49 | 56 | 49 | 52 |  |  |  |
| Ned | non-exposed | 29 | 35 | 35 | 33 | 342.88 | 1 | <0.0001 |
|  | virus exposed | 48 | 41 | 50 | 47 |  |  |  |

Analyses were conducted in software SPSS (version 19.0) by chi-square analysis.

**Figure 2.**
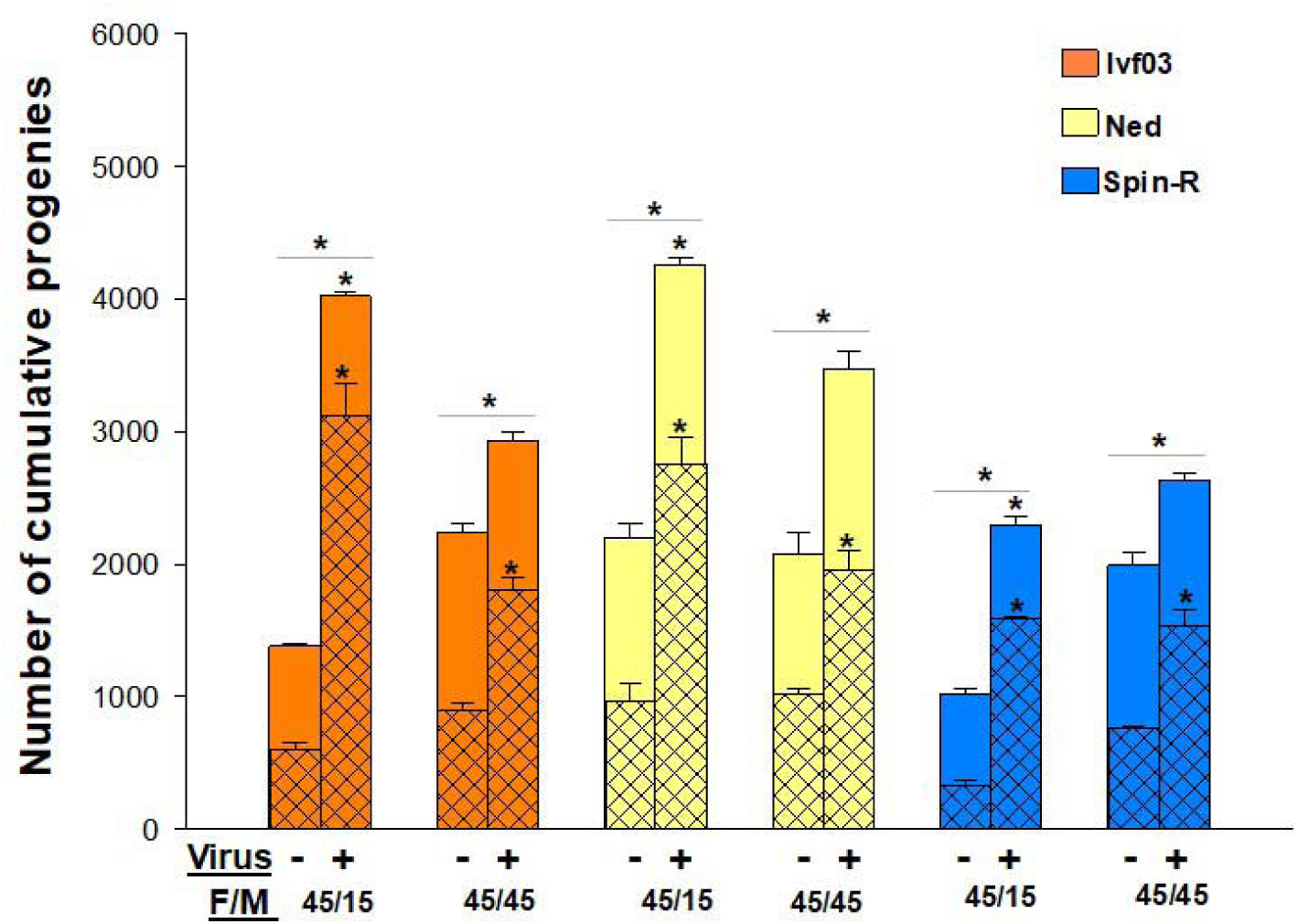
The offspring numbers of female (blank texture) and male (grid texture) in Ivf03, Spin-R, and Ned strains of *F. occidentalis* as determined by the TSWV status (virus exposed or non-exposed) of the parental thrips (in groups of 45 females and 15/45 males). An asterisk (*) indicates a significant difference in the number of thrips produced by virus exposed vs. non-exposed parents (one-way ANOVA, *P* ≤ 0.05). Values are means (± SE) of three replicates. A minus sign (-) indicates non-exposed treatment and a plus sign (+) indicates virus exposed treatment. M represents male and F represents female.

The male offspring numbers of three TSWV-exposed treatments (treatments 2, 3 and 4) were significantly higher than those of the non-exposed treatment 1. There was no significant difference among the female offspring of all four treatments. There was no significant difference among the offspring number of three TSWV-exposed treatments, means that the number of male and female offspring was not related to which parent was exposed to TSWV (Fig. 3).

**Figure 3.**
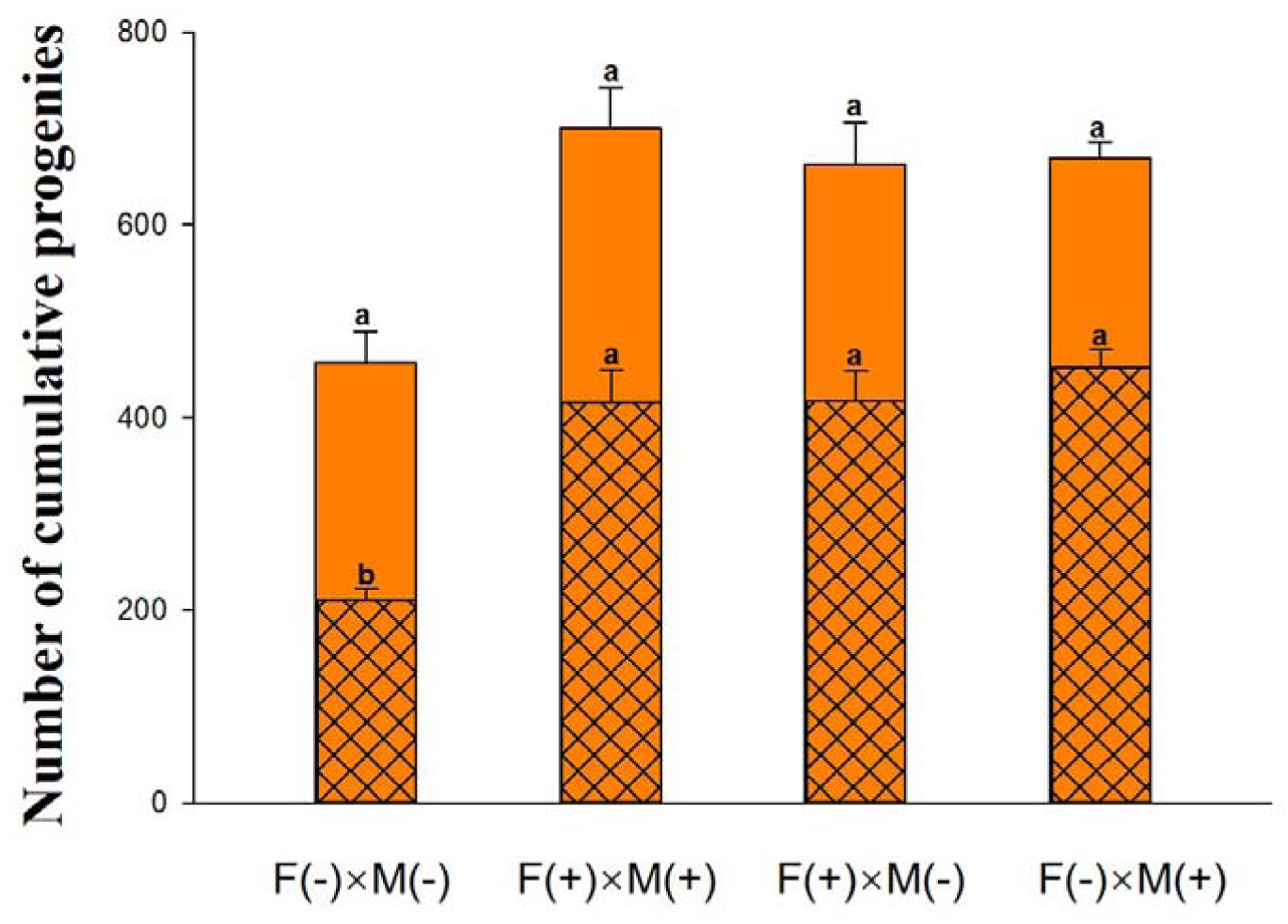
Numbers of offspring female (blank texture) and male (grid texture) of *F. occidentalis* in the four cross pair treatments (the parents were in groups of 10 females and 10 males). Different lowercase letters indicate significant differences (one-way ANOVA, *P* ≤ 0.05). Values are means (± SE) of three replicates. F and M indicate female and male thrips; (-) and (+) indicate TSWV-exposed and non-exposed thrips, respectively.

### 3.3 TSWV on the mating behaviour of *F. occidentalis*

During the mating, some behavioural aspects were more often observed in virus exposure treatments than in the non-exposed treatment (Fig. 4C, Suppl. Video 3), such as quick circling (Fig. 4D, E, F, Suppl. Video 1, 4, 5, 6), walking randomly and abdominal bending and swinging (Fig. 4D, Suppl. Video 4), female resistance by curving up abdomen (Fig. 4E, 4F, Suppl. Video 5, 6), female calming (Fig. 4D, E, Suppl. Video 4, 5), and female abdominal cleaning (Fig. 4F, Suppl. Video 6). Males exposed to TSWV (treatment 2 and 4) showed significantly higher male harassment rates than did non-exposed males (treatment 1 and 3; Table 4, Suppl. Video 2). There were no substantial differences in the pre-copulation period between treatments 1 and 2; however, a significantly shorter pre-copulation period in treatment 3 and a longer pre-copulation period in treatment 4 were observed (Fig. 4A). The copulation duration of TSWV-exposed treatments 2, 3 and 4 was significantly longer than that of non-exposed treatment 1 (Fig. 4B, Suppl. Video 3, 4, 5 and 6). During the first 1 h of video recording, female mating frequency was not significantly affected by TSWV in *F. occidentalis* (Table 4).

**Table 4.** Female mating frequency and male harassment rate (mean ± SE) of *F. occidentalis* in the four cross treatments

| Treatments | Female mating frequency | Male harassment rate |
| --- | --- | --- |
| F (-) × M (-) | 1.65±0.26a | 6.6±0.60b |
| F (+) × M (+) | 1.35±0.13a | 8.7±0.72a |
| F (+) × M (-) | 1.40±0.15a | 5.45±0.76b |
| F (-) × M (+) | 1.50±0.27a | 9.55±0.74a |

Different lowercase letters in a column indicate significant difference ( $P \leq 0.05$ ), according to one-way ANOVA . F and M indicate female and male thrips; (-) and (+) indicate TSWV-exposed and non-exposed thrips, respectively.

**Figure 4.**
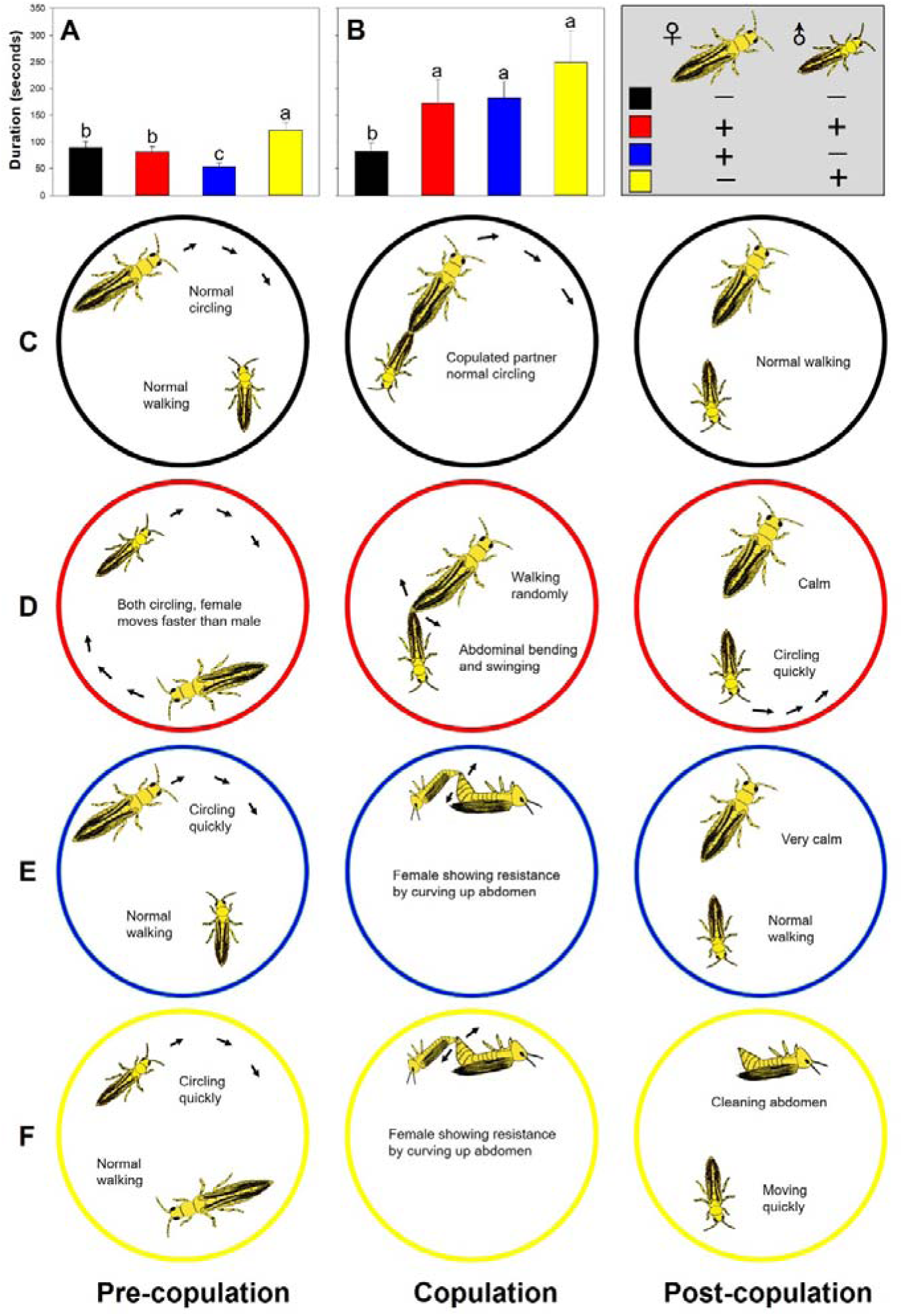
The duration of pre-copulation (A) and copulation (B), and schematic drawings of the most prominent posture of behaviours involved in mating (C, D, E, F) of male and female of *F. occidentalis* in the four cross pair treatments. Statistical significance in A and B (one-way ANOVA, *P* ≤ 0.05) is indicated by different lowercase letters. (-) and (+) indicate TSWV-uninfected and TSWV-infected thrips, respectively.

Infection with TSWV significantly reduced the accumulated re-mating frequency (P≤0.05; Fig. 5). Throughout the entire 20-day experimental period, 60% of the non-exposed females re-mated with non-exposed males (treatment 1), while in the TSWV-exposed treatments, only 25-30% of females re-mated with males (treatments 2, 3 and 4).

**Figure 5.**
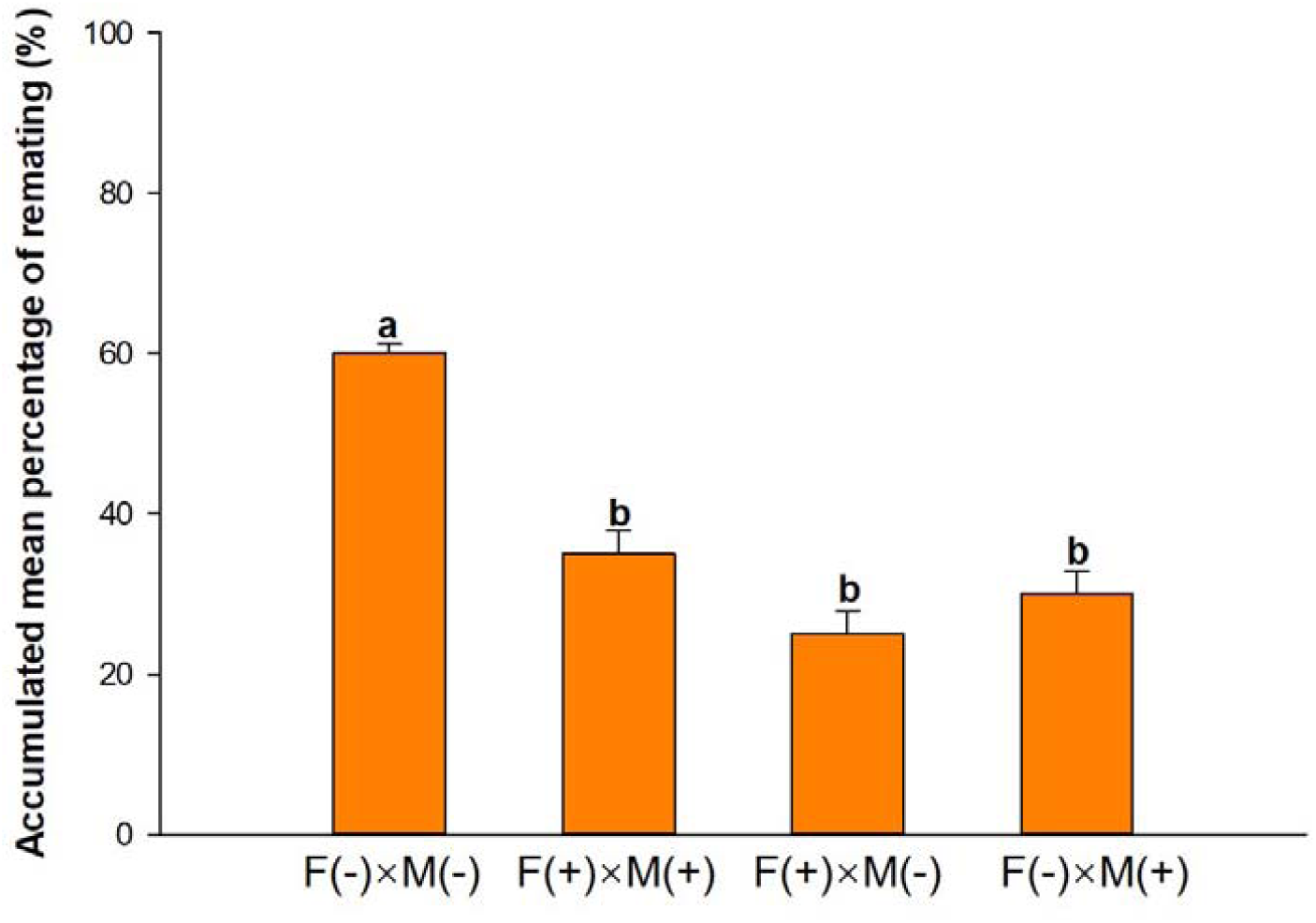
Accumulated percentage of re-mating *F. occidentalis* females in the four treatments. Statistical significance (chi-square analysis, *P* ≤ 0.05) is indicated by different lowercase letters. F and M indicate female and male thrips; (-) and (+) indicate TSWV-infected and uninfected thrips, respectively.

TSWV infection did not significantly affect the number of female progeny in any of the four treatments (Fig. 6). However, TSWV infection significantly increased the number of male offspring per female in all pairings in which at least one parent was infected with TSWV (treatments 2, 3, and 4) compared to uninfected thrips (treatment 1).

**Figure 6.**
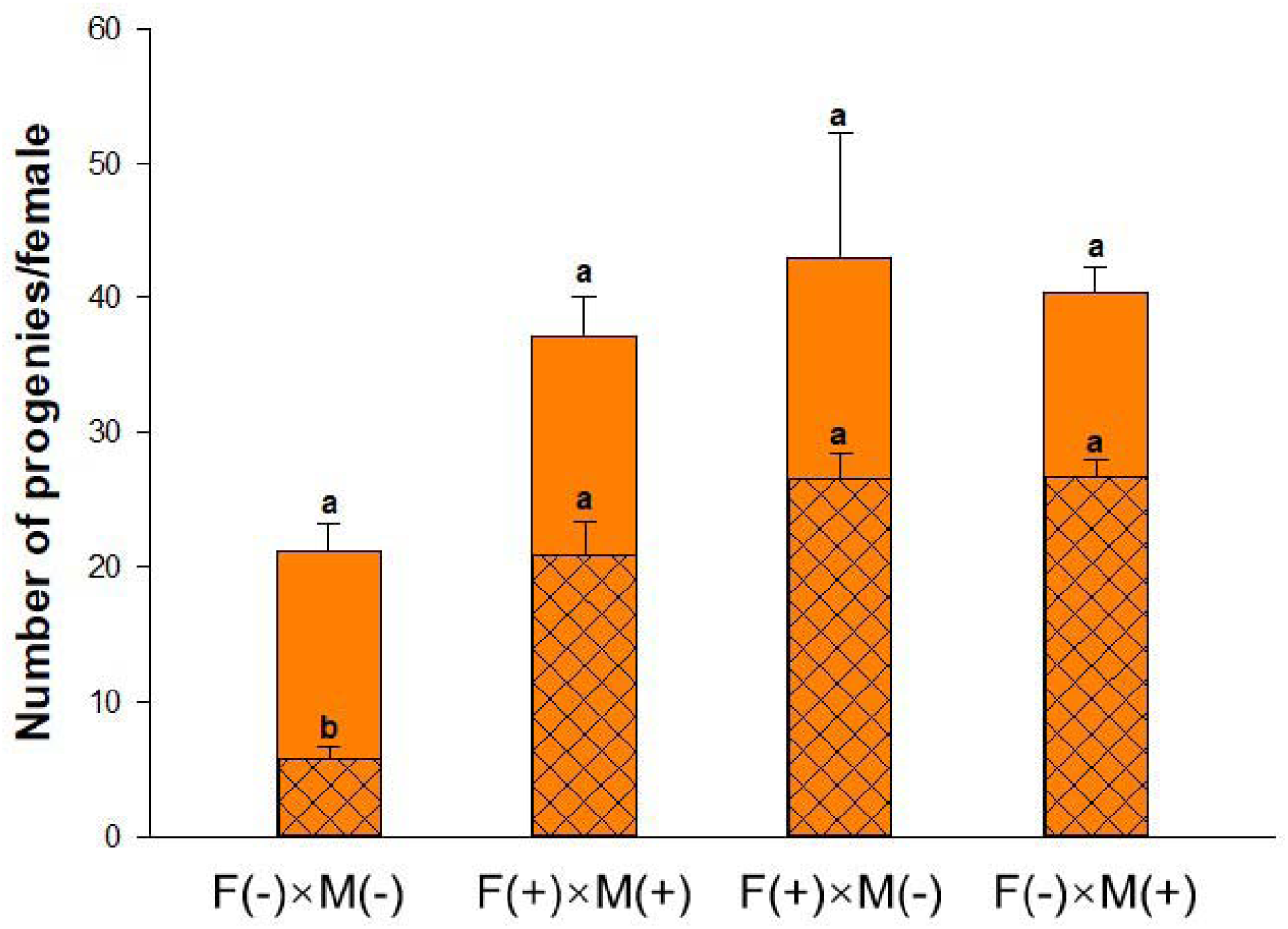
Mean numbers of offspring female (blank texture) and male (grid texture) per mated female of *F. occidentalis* in the four cross pair treatments. Values are means (± SE) (n=20). Different lowercase letters indicate significant differences (one-way ANOVA, *P* ≤ 0.05). F and M indicate female and male thrips; (-) and (+) indicate TSWV-uninfected and TSWV-infected thrips, respectively.

## 4 DISCUSSION

Plant viruses can alter the reproductive physiology and behaviour of the vector to facilitate viral transmission.^33,34^ In this study, TSWV manipulates the development and mating behaviour of its vector, *F. occidentalis*, to facilitate its transmission.

Specifically, when thrips were exposed to TSWV, their pre-adult period was shortened, and the adult longevity was prolonged, which is consistent with previous results^11,35^. The adult stage is the most effective TSWV transmission stage in thrips.^17^ Therefore, TSWV can modulate the developmental time of its vector to promote virus transmission.

The fecundity of both thrips parents exposed to TSWV was significantly higher (approximately 2-fold) than that of the non-exposed parents (Table 1). This result is consistent with Wijkamp et al.^25^, in which the net reproduction was 1.3-fold higher for virus-exposed than non-exposed thrips. Similarly, the fecundity of *F. occidentalis* was higher in TSWV-infected chickweed^36^ and tomato plants^27^, and TZSW-infected tomato plants.^35^ However, Ogada et al. (2013)^37^ reported a negative effect of TSWV on *F. occidentalis* fecundity. The difference may come from different strains of *F. occidentalis* and TSWV, and different host plants and methods used. Ogada et al (2013) used virgin female and observed the fecundity of arrhenotoky. In our study, the females were paired with males and both sexual and parthenogenesis reproduction were included. In addition, our cross-pair treatment indicated that as long as one parent was exposed to TSWV, the number of offspring will significantly increase. The increase in fecundity could be due to a nutritional gain in TSWV-infected plants compared to that in non-infected plants at nymph stage^8^ or female adult stage^37^. Selman et al.^38^ reported that TSWV-infected tomato plants increased total amino acids by 200-300%. Usually, higher concentrations of individual and total amino acids in infected foliage have a positive correlation with increased fecundity of its insect vector.^39,40^ In this study, thrips that became viruliferous by TSWV were fed TSWV-infected *D. stramonium* in the pre-adult stage but, like non-viruliferous thrips, were fed healthy bean pods in the adult stage. This suggests that improved nymph nutrition may explain the increased fecundity of virus-exposed thrips in our study. The copulation duration was longer for both female and male thrips infected with TSWV (Fig. 4B). Copulation and the nutrition from seminal fluid can stimulate females to oviposit more eggs.^41-43^ The increased copulation and prolonged longevity of females are other possible reasons for the increased fecundity of TSWV-exposed thrips.

In the current study, the non-exposed females of the three different strains of *F. occidentalis* produced more females than males, independent of the parental sex ratio (3:1 or 1:1) and mating pattern (in one pair or paired in groups). A female-biased sex ratio for *F. occidentalis* in nature was reported in previous studies. ^7,8^ Yuan et al.^44^ found that the filial sex ratio of *F. occidentalis* was 2-3:1 (female: male) for parental ratios of 1:1, 4:1, or 1:4. It is possible that if the external living conditions are stable, *F. occidentalis* can attain a stable sex ratio regardless of the preliminary sex ratio. According to the present results, this stable sex ratio is disturbed by the acquisition of TSWV. In this study, virulent females or females mated with virulent males produced significantly more progeny than virus-free females. More importantly, such increase in progeny was predominantly males, which led to a shift in the progeny sex ratio. Specifically, the female:male ratio was switched from 2-7:1 of the virulent parents to 0.6-1.1:1 of the virus-free parents. Given that 1) virus-free thrips prefer infected plants to feed, mate, and lay eggs^45,46^ and the mobility of the hatched nymphs is limited; and 2) virulent thrips prefer healthy plants, the resultant progenies are highly likely to acquire virus onsite through feeding. Previous findings showed that *F. occidentalis* males move much more between successive probes^22^ and are more efficient at transmitting virus than females^47^, the shift in the progeny sex ratio from female to male-biased apparently benefit the virus transmission.

Sex ratio distortion is expected to be especially prevalent in haplo-diploid species in part because the female parent has to control sex allocation.^48-50^ A recent study indicated that *Pteromalus puparum* negative-strand RNA virus 1 (PpNSRV-1) transmitted by both male and female wasps to offspring can also shift the sex ratio of the host wasp towards males but does so by decreasing the total number of female offspring because of the female killing element.^28^ However, the increase in male offspring leads to the distorted sex allocation of TSWV-exposed thrips in our study. It is generally accepted that *F. occidentalis* can re-mate five days after the first mating.^6^ However, TSWV infection appears to cause the females to refuse to mate again, and the frequency of re-mating decreased by approximately 50% (Fig. 5). For insects with multiple mating behaviour, females mate multiply can replenish depleted sperm or replace poor-quality sperm.^51^ Decreased remating may induce insufficient sperm acquired by female thrips. Thus, more unfertilized eggs develop into males, resulting in male-biased offspring.

Our group pairing experiment indicates that the parental sex ratio influences the offspring number. Both the female and male offspring numbers of 1:1 (45 females and 45 males) parents were significantly higher than those of 3:1 (45 females and 15 males) parents. In the 1:1 parent treatment, more males provided more mating opportunities for females than in the 3:1 treatment, copulation and the nutrition from seminal fluid can stimulate females to oviposit more eggs^41-43^, thus, more offspring were produced in the experimental conditions with enough food and shelter. Moreover, different parental sex ratios had different responses to TSWV infection. In the 3:1 treatment group, both male and female progenies were increased when exposed to TSWV. In the 1:1 treatment, however, only the male progeny increased significantly. Infection with TSWV caused males to become active, and they displayed more circular movements around the wall of the chamber than did healthy (uninfected) males (Fig. 4F, Suppl. Video 1), which led to higher rates of male harassment towards females for mating. There were three times the number of males in the 1:1 treatment than in the 3:1 treatment, and too much harassment from TSWV-infected males may disturb the oviposition of females, resulting in a reduced number of eggs.^52-54^

The three strains, Ivf03, Spin-R and Ned, used in the present study had almost the same response to TSWV exposure, although the fecundity of Spin-R was lower than those of the other two strains. Spin-R has an extremely high level of resistance to spinosad, and significant fitness cost was found in spinosad-resistant *F. occidentalis*.^55^ According to our present study, TSWV exposure can recover the reproductive fitness cost and improve the thrips population number. Meanwhile, the TSWV transmission efficiency of spinosad-resistant thrips was not different from that of normal thrips. ^23^ Considering the prevalence of *F. occidentalis* resistance to spinosad in the field,^4^ more research needs to investigate the interaction between pesticide-resistant thrips and TSWV.

Understanding the interaction between plant pathogens and their vectors is important for successful pest control. The transmission of insect-borne plant viruses depends on the abundance and behaviour of their vectors.^26^ TSWV manipulates the development time, mating behaviour, fecundity and sex allocation of its vector *F. occidentalis* to enhance its spread. Consequently, for the long-term, sustainable management of TSWV and thrips, a monitoring program of earlier detection of the virus in plants and thrips by using DAS-ELISA, RT-PCR or virus detection strip must be implemented in the field. And a reduced economic threshold for vector (thrips) control should be in consideration in areas where the virus is already present.

## ACKNOWLEDGEMENTS

This study was funded by the National Key R&D Program of China (2017YFD0201000), Natural Science Foundation of China (31772199, 31572037), the Science and Technology Innovation Program of the CAAS (AAS-ASTIP-IVFCAAS), Beijing Leafy Vegetables Innovation Team of Modern Agro-industry Technology Research System (BAIC07-2018), and Beijing Key Laboratory for Pest Control and Sustainable Cultivation of Vegetables.

