## Supplementary material for "*Tomato spotted wilt virus* manipulates the reproduction of its insect vector, western flower thrips (*Frankliniella occidentalis*), to facilitate transmission": Suppl. Video: Supporting Videos details information.docx

**Details information about Supplementary materials 1, 2, 3, 4, 5 and 6 videos.**

**Video S1**

The video shows the circular movement of a TSWV infected male in a Petri dish with a non-infected female during pairing. The male is very active and makes circular movement before copulation.

**Video S2**

The video shows a male harassing a female thrips. The male is TSWV infected and the female is not. Female shows resistance to mate despite male’s attempts.

**Video S3**

The video shows the pre-copulation and copulation behaviour of a TSWV uninfected male and female couple of *F.occidentalis* during mating. Female thrips willing to mate with male thrips, more active, and after mating move fast.

**Video S4**

The video shows the pre-copulation and copulation behaviour of a male and female couple of *F.occidentalis* in which both sexes were TSWV infected. Both male and female thrips are very active. Female thrips is reluctant to mate.

**Video S5**

The video shows the pre-copulation and copulation behaviour in which the female is infected with TSWV but the male is uninfected. Female thrips makes circular position in pre-copulation to avoid mating with male. It shows highly resistance for mating. Also, female thrips observed very calm during and after mating. Copulation duration is higher in this treatment.

**Video S6**

The video shows the pre-copulation and copulation behaviour in which the male is TSWV infected and the female is uninfected. Male becomes more active and female is calm during mating. Female also shows resistance for virus infected male. After mating, female uplifts its abdomen and looks rubbing its body. It is unusual behaviour among all treatments. After mating, male attempts again for re-mating.
